# Occurrence and associated characteristics of a mutated *ant(6’)-Ia* gene among *Enterococcus faecium* strains expressing phenotypic susceptibility to high levels of streptomycin

**DOI:** 10.1101/2020.12.28.424548

**Authors:** Stephanie Silva Rodrigues Souza, Adriana Rocha Faria, Andréa Andrade Rangel Freitas, Paul J Planet, Vânia Lúcia Carreira Merquior, Lúcia Martins Teixeira

## Abstract

Enterococcal high-level resistance to streptomycin (HLR-S) (MIC ≥ 2000 µg/ml), conferred by production of a variety of aminoglycoside modifying enzymes (AMES), has been reported worldwide and a nucleotidyltransferase (ANT) enzyme encoded by the *ant(6’)-Ia* gene is frequently associated with this phenotype. However, during a study conducted by our group on whole genome sequencing (WGS) analyses of *Enterococcus faecium* isolates, we observed that 32 *E. faecium* strains identified as susceptible to high-levels of streptomycin by the disk diffusion method had the of *ant(6’)-Ia* gene annotated in their genomes. Antimicrobial susceptibility to streptomycin was reassessed by phenotypic testing and the presence of the *ant(6’)-Ia* gene was confirmed by PCR in all the isolates. Alignment of the *ant(6’)-Ia* gene with a reference sequence revealed a deletion of the first 48 nucleotides and four nonsynonymous mutations, leading to the substitution of a Glutamine to Methionine and an Aspartic Acid to Asparagine in the amino acid sequence. The protein structure was modelled by using the Phyre2 platform and the results indicated alterations in the N-terminus region leading to changes in the predicted binding site. Also, by searching the NCBI database we identified the genomes of 71 strains carrying the mutated gene. MLST analysis revealed that most strains carrying the mutated gene, including those described in this study belonged to hospital-adapted lineages, suggesting the occurrence of clonal dissemination of a subset of mutated isolates.

**HIGHLIGHTS:**

- The presence of a mutated *ant(6’)-Ia* gene was identified among *Enterococcus faecium* isolates expressing phenotypic susceptibility to high levels of streptomycin.
- Nonsynonymous mutations and inactivating changes in the *ant(6’)-Ia* gene led to incongruities between phenotypes and genotypes.
- Alterations in the amino acid sequence had impacts on protein structure, with changes in the N-terminus region and the binding site.

## 1. Introduction

Enterococci are intrinsically resistant to low levels of aminoglycosides due to reduced membrane permeability. Consequently, the synergistic combination of an aminoglycoside with a cell wall active agent such as a beta-lactam or a glycopeptide is recommended as the treatment of choice for critical enterococcal infections, such as bacteremia and endocarditis [1,2]. Nevertheless, the efficacy of the above therapeutic combination has been impaired since the emergence and dissemination of enterococcal strains showing high-level resistance to aminoglycosides (HLR-A), leading to MICs ≥ 2000 μg/mL, and posing significant limitations in treatment options [3].

Streptomycin is an alternative for treatment of infections caused by strains showing high-level resistance to gentamicin (HLR-G), considering that the most frequent resistance mechanisms also eliminate the activity against all other clinically relevant aminoglycosides, including amikacin, kanamycin, netilmicin, tobramycin and sisomicin. However, the occurrence and spread of strains with high-level resistance to streptomycin (HLR-S) has been reported around the world since the early 1970s [4-6].

HLR-S is commonly due to point mutations in the ribosomal proteins or to the acquisition of genes encoding aminoglycoside-modifying enzymes (AMEs) [7]. In diagnostic laboratories, the disk diffusion method using 300 μg streptomycin disks is the standard procedure for detecting the HLR-S phenotype among enterococci [8,9]. Additionally, PCR screening for detecting resistance determinants is a useful and convenient molecular tool when the strain is categorized in the breakpoint range considered inconclusive (7 to 9 mm).

Among enterococci, the most common AME associated with the inactivation of streptomycin is a nucleotidyltransferase encoded by the *ant(6)-Ia* gene, also recognized as the *aadE* gene [10]. This gene is present in the gene cluster *aadE-sat4-aph3* which is generally integrated into the transposable element Tn*5405*-like, disseminated among *E. faecium* of different ecological origins as well as *Staphylococcus aureus* and coagulase-negative staphylococci [11-13].

In *Enterococcus faecium*, the dissemination of HLR-S strains and respective genetic determinants is generally associated with hospital-adapted lineages that have been reported as causes of outbreaks worldwide [14-18].

The aim of this study was to investigate the unexpected presence of the *ant(6)-Ia* gene and related characteristics among *Enterococcus faecium* strains expressing phenotypic susceptibility to high levels of streptomycin.

## 2. Materials and Methods

### 2.1 Bacterial strains

A total of 32 *Enterococcus faecium* isolates presenting incongruent results derived from phenotypic and genotypic evaluation of susceptibility to streptomycin were included in the present study. They were identified during previous studies conducted by our group on whole-genome sequencing (WGS) analyses of *Enterococcus faecium*. The isolates were recovered from patients in 18 different health institutions of Rio de Janeiro, Southeastern region of Brazil, over a period of nine years (2008 to 2017). As their isolation took place as part of the standard patient care procedures, ethical approval for their use was not required.

Early identification of the isolates, based on physiological tests [19], was confirmed by matrix assisted laser desorption ionization-time of flight mass spectrometry (MALDI-TOF MS) analysis (Microflex LT MALDI-TOF MS system; Bruker Daltonics), following the instructions of the manufacturer. For subsequent tests, bacterial cells were grown on blood agar plates (Trypticase soy agar supplemented with 5% defibrinated sheep blood; Plast Labor) at 36°C± 1°C for 18 to 24 h.

### 2.2 Antimicrobial susceptibility testing

Susceptibility to streptomycin was evaluated by the disk-diffusion method, using high-level streptomycin (300 μg) disks (Oxoid Ltd), according to the Clinical and Laboratory Standards Institute recommendations [9]. Minimum inhibitory concentrations (MICs) were determined by an agar dilution method using Mueller-Hinton agar (Becton, Dickinson and Company) supplemented with streptomycin (Sigma-Aldrich Co.) in concentrations ranging from 0.25 to 4,096 mg/L). Interpretation of the results was based on the CLSI guidelines [9].

### 2.3 Detection of the ant(6’)-Ia gene

The presence of the *ant(6’)-Ia* gene was evaluated by PCR assays according to Swenson et al., [8]. Bacterial DNA was extracted using the Chelex 100 (Bio-Rad, Hercules, CA, USA) resin [20]. The reference strains *E. faecalis* ATCC49533 and *E. faecalis* ATCC29212 were used as positive and negative controls, respectively. The PRIMER-blast tool was also used to detect the specific annealing regions [21].

### 2.4 Whole-Genome Sequencing analysis (WGS)

A single bacterial colony grown on a blood agar plate was inoculated into 5□mL of tryptic soy broth and incubated overnight at 37°C. Genomic DNA was extracted from 1.5□mL of that culture by using a Wizard genomic DNA purification kit (Promega, Madison, WI, USA), prepped using the Nextera XT kit, and sequenced on a HiSeq 2500 sequencer (Illumina Inc., San Diego, CA, USA) with 125-pb paired-end reads. Trimming of the reads was accomplished by using the Trim Galore v. 0.6.2 (https://www.bioinformatics.babraham.ac.uk/projects/trim_galore/) software and the quality metrics was assessed using FastQC [22]. High-quality reads were assembled and annotated using resources of the Pathosystems Resource Integration Center (PATRIC 3.4.11) [23]. Moreover, the identification of acquired antibiotic resistance genes was also evaluated using Resfinder [24].

The fasta files of amino acid and nucleotide sequences from *ant(6’)-Ia* gene were extracted and aligned with the reference sequence of the gene cluster *aadE-sat4-aph3* (accession no. AF330699.1) using the MEGA software (version 7), and also, evaluated against the NCBI database using BLAST analysis.

The genome sequences reported in this work have been deposited at DDBJ/ENA/GenBank under the BioProject accession number PRJNA646532.

### 2.5 Protein modeling

The amino acid sequences were used to predict the 3D protein structure and binding sites in the PHYRE2 server and 3DLigandSite, respectively [25,26]. The protein display and images production were made by using EzMol 1.3 [27].

## 3. Results

In the present study, the occurrence of the incongruity between phenotypes and genotypes of resistance to streptomycin was investigated in 32 *E. faecium* strains that carried the *ant(6’)-Ia* gene as detected after annotation of the whole-genome. These strains were previously classified as susceptible to high-levels of streptomycin by the disk-diffusion method and results were confirmed by repeating the tests (Table 1).

**TABLE 1.**
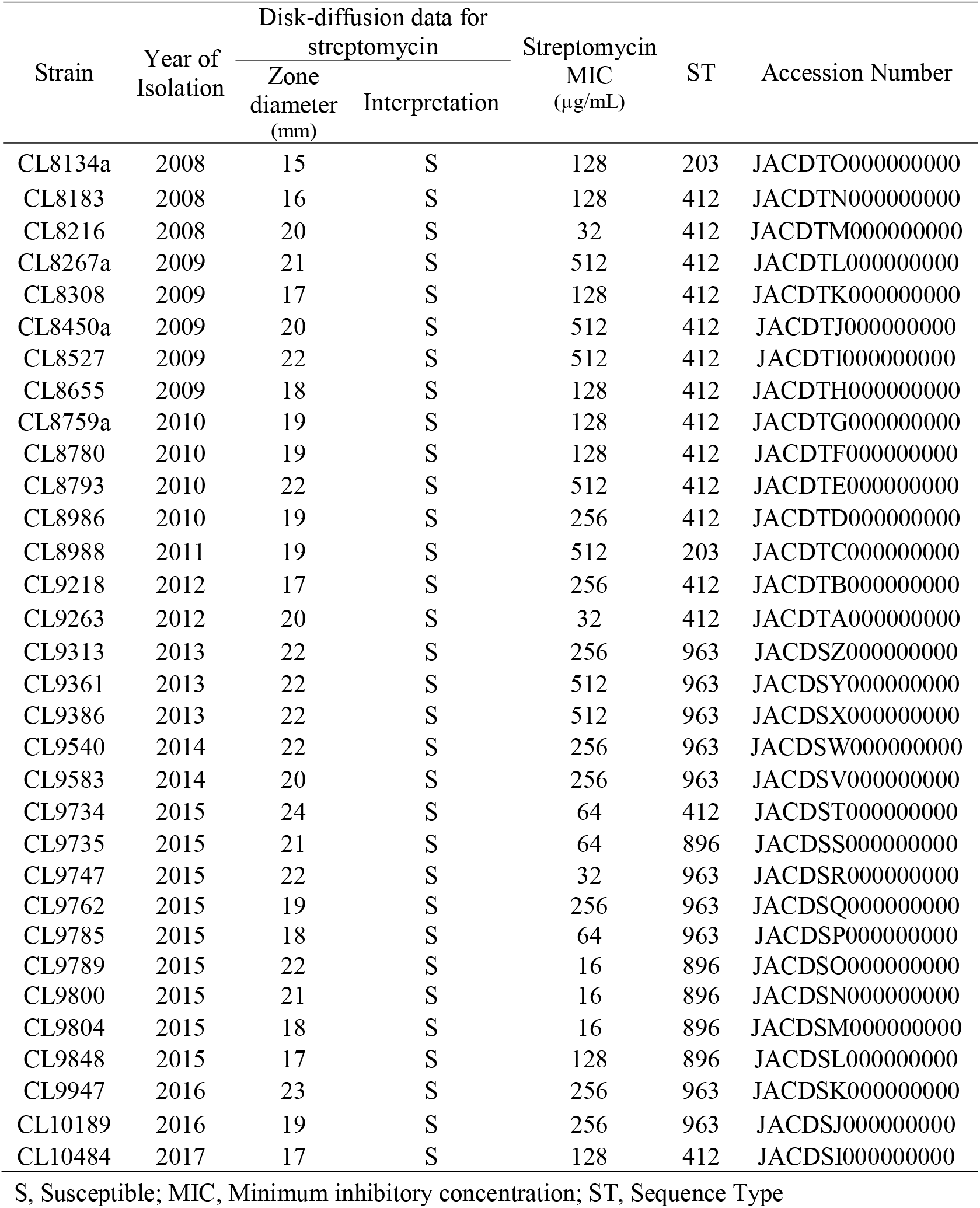
Characteristics of the thirty-two *Enterococccus faecium* isolates possessing a mutated *ant(6’)-Ia* gene.

In order to confirm the consistency of these findings, determination of streptomycin MICs and PCR-based detection of the *ant(6’)-Ia* gene were performed. The MIC values obtained ranged from 16 to 512 μg/mL (Table 1), confirming that all 32 isolates had phenotypes indicative of susceptibility to high-levels of streptomycin. Moreover, PCR product for *ant(6’)-Ia* was also detected in all strains. The strains were classified as four distinct sequence types (STs) (Table 1). Moreover, by searching the NCBI database, 71 additional *E. faecium* strains carrying the *ant(6’)-Ia* gene with the exact same changes were identified (Table S1). These strains belonged to eight different STs, with a predominance of ST736.

By WGS analysis, a deletion of the first 48 nucleotides, leading to changes in the amino acid sequence was identified in the *ant(6’)-Ia* gene. Also, four nonsynonymous mutations in the positions 49 (C-A), 50 (A-T), 52 (G-A) and 54 (T-C) were identified in all strains. These point mutations led to the substitution of a Glutamine to Methionine and an Aspartic Acid to an Asparagine (Figure 1)

**FIGURE 1.**
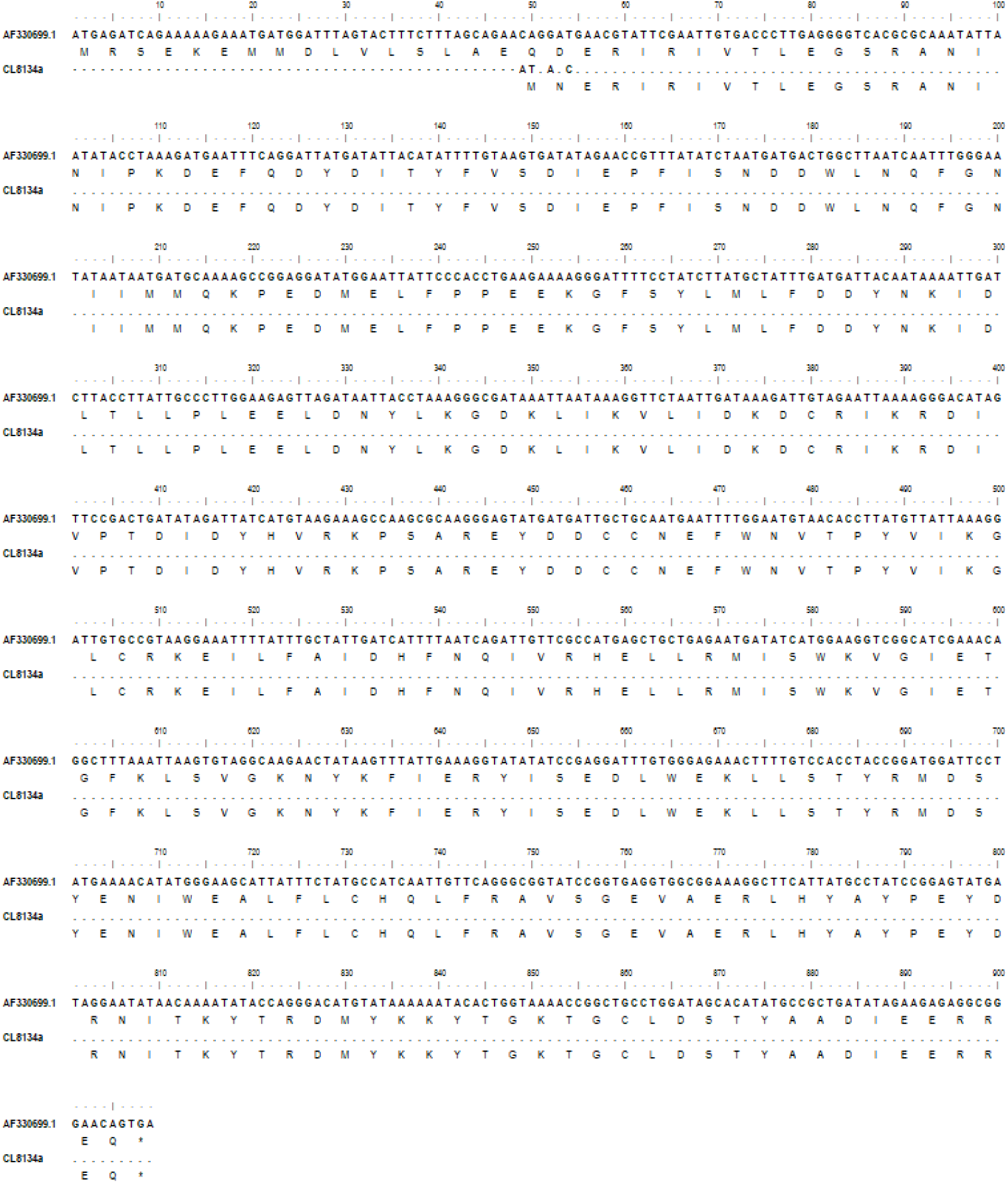
Sequences of nucleotide and amino acid obtained from *ant(6’)-Ia* gene of *Enterococcus faecium* CL8134a, an incongruent strain identified in the presente study, and the reference strain (AF330699.1), aligned in the MEGA software using CLUSTALW.

The 3D model of the protein was predicted by PHYRE2 with a 100% of confidence and 85% of coverage. The predicted skeleton structure of ANT(6’)-IA showed alterations in the N-terminus region when compared with the reference model (Figure 2A; 2B). Also, these alterations in the protein structure led to changes in the predicted binding site (Fig 2C; 2D) and consequently loss of the protein function. The mutated ANT(6’)-IA protein had nine amino acid residues predicted as binding sites.

**FIGURE 2.**
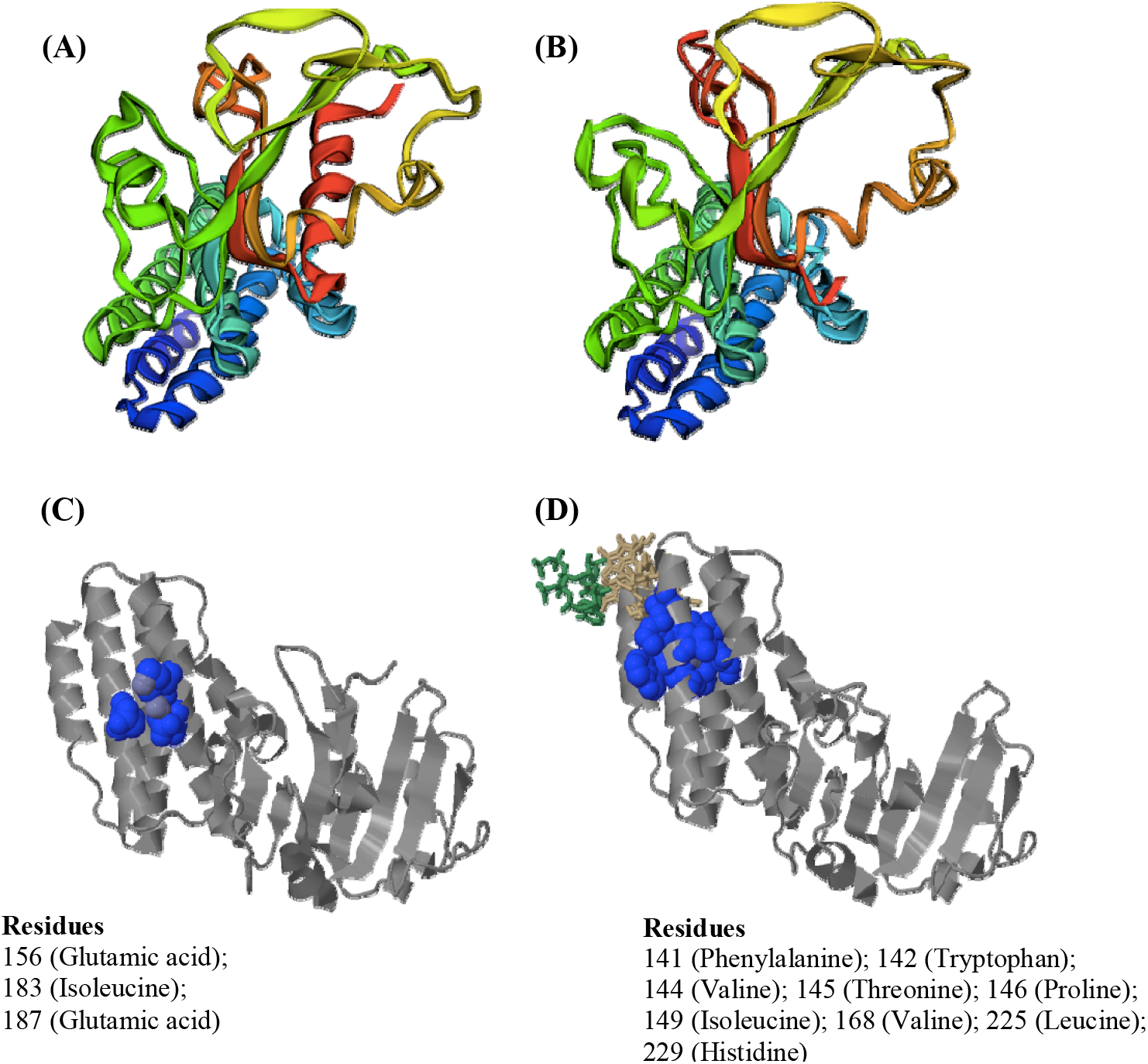
Structural model of ANT(6’)-IA protein and predicted biding sites. 3D protein models colored by rainbow N→C terminus of the reference sequence (AF330699.1) **(A)** and the mutated sequence **(B)**. Predicted binding sites (in blue) and the respective amino acid residues of the reference sequence (AF330699.1) **(C)** and the mutated sequence **(D)**.

## 4. Discussion

Surveillance of HLR-A and respective genetic bases has a significant clinical impact, not only for the choice of appropriate therapy but also for understanding the mechanisms and epidemiology of this type of resistance. The potential provided by genotype-based technology, such as PCR and more recently by WGS, for fast and accurate detection has led to an increase in the use of these methodologies to predict antibiotic resistance in routine laboratories [28,29]. However, the occurrence of discrepancies between results of phenotypic and genetic testing is still poorly documented and understood.

A recent report from the EUCAST subcommittee gave a baseline document for further discussion on the use of WGS as a tool to infer antimicrobial susceptibility, once there is insufficient evidence to support this practice to guide clinical decisions [30]. In fact, if the predictions of streptomycin susceptibility of the 32 isolates included in the present study were only based on genotype screening, they would be considered resistant. This finding was only called to our attention because we had previously tested these isolates for phenotypic resistance. Moreover, the confirmation of the presence of antimicrobial resistance genes by PCR opens an important discussion about the prediction of antibiotic resistance based in molecular techniques only, in the absence of correlated MIC data. Although molecular techniques are useful tools, they also have intrinsic limitations. In PCR assays nucleotide insertions or deletions in regions outside the PCR product that may inactivate the gene would not be detected, giving rise to false positive results. In our case, using the PRIMER-blast tool, we have observed that the primer pair routinely used for *ant(6’)-Ia* detection has its annealing region outside the area where deletions and mutations are located.

Inconsistencies between phenotypic and genotypic evaluation of susceptibility to streptomycin have been reported occasionally. Hospital-associated enterococcal strains carrying the *aadE* gene with a non-HLSR phenotype were described by Leelaporn et al. [31]. Considering nonhuman sources, isolates containing *ant(6’)-Ia* but susceptible to streptomycin were also observed among enterococci recovered from poultry carcasses [32]. Furthermore, in more recent studies using WGS to predict antibiotic resistance in *Escherichia coli* and *Salmonella*, high degrees of phenotype and genotype incongruences related to aminoglycoside susceptibility have been reported [33-35].

Bacterial strains carrying non-functional genes seem to be a relatively common occurrence, and the genetic bases that led to these phenomena are diverse. The presence of premature stop codons or a non-functional promoter has been described [36,37]. Genetic rearrangements in the transposon carrying the *ant(6’)-Ia* gene also could be one explanation for the deletions and mutations identified in this study. In fact, the rearrangements in the cluster *aadE-sat4-aph3* and the presence of IS elements inside the Tn*5405*-like also have been demonstrated [38].

The presence of mutated *ant(6’)-Ia* gene in a large number of strains not only in our country, but also in the USA, as we observed in genomes deposited at the NCBI database (Supplementary data), may indicate a unexpectedly wide dissemination of this non-functional gene. Interestingly, the majority of the strains possessing the mutated gene belonged to hospital-adapted lineages, suggesting the occurrence of clonal dissemination.

A common feature of resistance that is acquired by HGT is the co-selection of resistance markers [39]. Such selection could explain the maintenance of a non-functional *ant(6’)-Ia* gene. Indeed, there is a strong association between the presence of the gene cluster *aadE-sat4-aph3* and the *ermB* (erythromycin resistance) gene in Tn*5405* [13, 38]. Moreover, this transposon is known to be harbored on different conjugative multiresistance plasmids that also carried *Tn*1546-like transposon, which is associated with the circulation of the *vanA* gene, the major determinant responsible for the spread of vancomycin resistance among *Enterococcus* spp. [10].

The genetic alterations in the amino acid sequence caused by deletions and mutations led to changes in the N-terminal domain of the protein structure. A recent study with another adenylyltransferase enzyme, ANT(2”)-Ia, revealed that the N-terminal domain is conserved among other members in the protein family, including ANT (4’)-Ia, ANT (4”)-Ib and ANT(6)-Ia. Moreover, the authors demonstrated that the catalytic site (Asp86) of the ANT(2”)-Ia enzyme is located in this domain, and showed that any substitution led to the complete loss of enzyme activity [40].

In conclusion, we have found that *E. faecium* strains showing susceptibility to high levels of streptomycin can carry a mutated *ant(6’)-Ia* gene due to a deletion of the first 48 nucleotides and four nonsynonymous mutations, leading to changes in the amino acid sequence of the predicted binding site of the enzyme and the consequent loss of function. Although molecular techniques are important tools to detect and identify resistance to antimicrobials, they may not always correctly predict function or expression. Phenotypic tests may still remain an important tool to guide the choice of antibiotic therapy and their systematic use in combination with genotypic approaches could help to clarify the impact of incongruent phenotypes and genotypes.

## Acknowledgments

This work was supported in part by Coordenação de Aperfeiçoamento de Pessoal de Nível Superior (CAPES)-Finance Code 001, Instituto Nacional de Pesquisa em Resistência Antimicrobiana (INPRA), Conselho Nacional de Desenvolvimento Científico e Tecnológico (CNPq), Fundação de Amparo à Pesquisa do Estado do Rio de Janeiro (FAPERJ), Brazil, and Columbia University Global Centers/FAPERJ Collaborative Grant and the Columbia University President’s Global Innovation Fund.

