## Supplementary data for "Occurrence and associated characteristics of a mutated *ant(6’)-Ia* gene among *Enterococcus faecium* strains expressing phenotypic susceptibility to high levels of streptomycin"

| Strain | Accession number | Country-Year | Sequence Type (ST) |
|---|---|---|---|
| E232 | GCA_002777275.1 | USA - 2013 | ST736 |
| E240 | GCA_002761255.1 | USA - 2013 | ST736 |
| E243 | GCA_002761275.1 | USA - 2013 | ST736 |
| ER04484.3A | GCA_002848665.1 | USA - 2015 | ST736 |
| ER04526.5A | GCA_002848725.1 | USA - 2015 | ST736 |
| ER04462.3A | GCA_002848685.1 | USA - 2015 | ST736 |
| ER04120.3A | GCA_002848645.1 | USA - 2015 | ST736 |
| UAMSEF_47 | GCA_004300625.1 | USA - 2018 | ST412 |
| UAMSEF_21 | GCA_004300315.1 | USA - 2018 | ST412 |
| UAMSEF_29 | GCA_004300195.1 | USA - 2018 | ST736 |
| UAMSEF_28 | GCA_004300355.1 | USA - 2018 | ST736 |
| UAMSEF_44 | GCA_004299885.1 | USA - 2018 | ST736 |
| UAMSEF_33 | GCA_004299975.1 | USA - 2018 | ST736 |
| UAMSEF_32 | GCA_004299965.1 | USA - 2018 | ST203 |
| UAMSEF_45 | GCA_004299905.1 | USA - 2018 | ST736 |
| UAMSEF_25 | GCA_004300375.1 | USA - 2018 | ST448 |
| UAMSEF_42 | GCA_004300115.1 | USA - 2018 | ST203 |
| UAMSEF_27 | GCA_004299875.1 | USA - 2018 | ST736 |
| EF_534 | GCA_004151795.1 | USA - 2018 | ST18 |
| EF_531 | GCA_004151925.1 | USA - 2018 | ST18 |
| EF_536 | GCA_004151785.1 | USA - 2018 | ST18 |
| EF_500 | GCA_004152605.1 | USA - 2018 | ST412 |
| EF_508 | GCA_004152205.1 | USA - 2018 | ST78 |
| EF_511 | GCA_004152175.1 | USA - 2018 | ST736 |
| EF_532 | GCA_004151875.1 | USA - 2018 | ST18 |
| AALTL | GCA_002880635.1 | USA - 2009 | ST736 |
| IHC30 | GCA_003350705.1 | USA - 2007 | ST412 |
| ET12_17 | GCA_002750905.2 | Brazil - 2017 | ST896 |
| ET13_17 | GCA_002751055.2 | Brazil - 2017 | ST896 |
| ET14_17 | GCA_002750945.2 | Brazil - 2017 | ST896 |
| 4278 | GCA_003242945.1 | Brazil - 2014 | ST896 |
| IHC22 | GCA_003335725.1 | USA - 2011 | ST736 |
| IHC23 | GCA_003335735.1 | USA - 2007 | ST412 |
| IHC33 | GCA_003323695.1 | USA - 2009 | ST896 |
| UEL170 | GCA_003697805.1 | Brazil - 2009 | ST412 |
| EF_539 | GCA_004151775.1 | USA – 2018 | ST18 |
| EF_512 | GCA_004152505.1 | USA – 2018 | ST78 |
| E39 | GCA_001635875.1 | USA – 2009 | ST736 |
| ISMMS_VRE_5 | GCA_001721025.1 | USA – 2011 | ST736 |
| ISMMS_VRE_7 | GCA_001721065.1 | USA - 2011 | ST17 |
| ISMMS_VRE_10 | GCA_001721105.1 | USA - 2011 | ST736 |
| ISMMS_VRE_6 | GCA_001953715.1 | USA - 2011 | ST736 |
| HMSC070F12 | GCA_001837605.1 | USA - 2016 | ST412 |
| LIM1590 | GCA_002005825.1 | Brazil - 2013 | ST963 |
| LIM918 | GCA_002005815.1 | Brazil - 2012 | ST896 |

| | | | |
|---|---|---|---|
| LIM4247 | GCA_002005885.1 | Brazil - 2014 | ST896 |
| LIM1759 | GCA_002005915.1 | Brazil - 2014 | ST896 |
| IHC8 | GCA_002158355.1 | USA – 2007 | ST412 |
| IHC13 | GCA_002158245.1 | USA – 2006 | ST412 |
| R499 | GCA_000294875.2 | USA - 2013 | ST412 |
| P1137 | GCA_000295135.2 | USA – 2013 | ST18 |
| P1140 | GCA_000295015.1 | USA – 2013 | ST412 |
| ERV168 | GCA_000295075.2 | USA – 2013 | ST412 |
| ERV165 | GCA_000295235.2 | USA – 2013 | ST412 |
| EnGen0314 | GCA_000394635.1 | USA - 2012 | ST736 |
| EnGen0319 | GCA_000394695.1 | USA - 2012 | ST736 |
| EnGen0323 | GCA_000394735.1 | USA - 2012 | ST736 |
| EnGen0312 | GCA_000394755.1 | USA - 2012 | ST736 |
| EnGen0376 | GCA_000407085.1 | USA - 2012 | ST736 |
| EnGen0375 | GCA_000407625.1 | USA - 2012 | ST736 |
| EnGen0377 | GCA_000407105.1 | USA - 2012 | ST736 |
| MRSN33034 | GCA_001481405.1 | USA – 2015 | ST78 |
| HMSC073E08 | GCA_001811715.1 | USA - 2016 | ST736 |
| HMSC073E07 | GCA_001810595.1 | USA - 2016 | ST736 |
| HMSC072G01 | GCA_001809135.1 | USA - 2016 | ST736 |
| HMSC076D08 | GCA_001815195.1 | USA - 2016 | ST412 |
| HMSC065H03 | GCA_001815195.1 | USA - 2016 | ST736 |
| HMSC060E05 | GCA_001814465.1 | USA - 2016 | ST18 |
| HMSC063C12 | GCA_001814315.1 | USA - 2016 | ST412 |
| HMSC065H12 | GCA_001813575.1 | USA - 2016 | ST412 |
| HMSC072F07 | GCA_001812654.1 | USA - 2016 | ST736 |

List of strains obtained from the NCBI database carrying a mutated *ant(6`)-Ia* gene and their respective characteristics.
